# A high-affinity antibody against the CSP N-terminal domain lacks *Plasmodium falciparum* inhibitory activity

**DOI:** 10.1101/2020.01.13.904425

**Authors:** Elaine Thai, Giulia Costa, Anna Weyrich, Rajagopal Murugan, David Oyen, Katherine Prieto, Alexandre Bosch, Angelo Valleriani, Nicholas C. Wu, Tossapol Pholcharee, Stephen W. Scally, Ian A. Wilson, Hedda Wardemann, Jean-Philippe Julien, Elena A. Levashina

**Author notes:** To whom correspondence should be addressed: Elena Levashina Vector Biology Unit Max Planck Institute for Infection Biology Campus Charité Mitte Charitéplatz 1 10117 Berlin, Germany Jean-Philippe Julien The Hospital for Sick Children Research Institute Peter Gilgan Centre for Research and Learning (PGCRL), Room 20-9703 686 Bay St. Toronto ON Canada M5G 0A4 Hedda Wardemann B Cell Immunology (D130) Deutsches Krebsforschungszentrum Im Neuenheimer Feld 280 69120 Heidelberg, Germany. These authors contributed equally (Elaine Thai, Giulia Costa, Anna Weyrich).

## Abstract

Malaria is a global health concern and research efforts are ongoing to develop a superior vaccine to RTS,S/AS01. To guide immunogen design, we seek a comprehensive understanding of the protective humoral response against *Plasmodium falciparum* circumsporozoite protein (PfCSP). In contrast to the well-studied responses to the repeat region and the C-terminus, the antibody response against the N-terminal domain of PfCSP (N-CSP) remains obscure. Here, we characterized the molecular recognition and functional efficacy of the N-CSP-specific monoclonal antibody 5D5. The crystal structure at 1.85 Å resolution revealed that 5D5 binds an *α*-helical epitope in N-CSP with high affinity through extensive shape and charge complementarity, and the unusual utilization of an N-linked glycan. Nevertheless, functional studies indicated low 5D5 binding to live Pf sporozoites, and lack of sporozoite inhibition *in vitro* and in mosquitoes. Overall, our data on low recognition and inhibition of sporozoites do not support the inclusion of the 5D5 epitope into the next generation of CSP-based vaccines.

**Summary Statement:** The *Plasmodium falciparum* sporozoite surface protein, PfCSP, is an attractive vaccine target, but the antibody response against the CSP N-terminal domain has remained understudied. Here, to guide immunogen design, Thai et al. provide insights into the binding motif and functional efficacy of the N-terminal domain-specific monoclonal antibody, 5D5.

## Introduction

Malaria is a vector-borne disease of global importance. In 2017, an estimated 219 million cases were reported, resulting in 435,000 deaths (WHO, 2018). The majority of deaths are caused by *Plasmodium falciparum* (Pf), making this parasite a central focus of research efforts for the development of effective therapeutic interventions. Anti-infection vaccines target the sporozoite stage of the Pf life cycle as parasites are transmitted to the human host by infected female *Anopheles* mosquitos during a blood meal. It has been established four decades ago that mAbs targeting the sporozoite surface circumsporozoite protein (CSP) are capable of neutralizing *Plasmodium* infection (Potocnjak et al., 1980; Yoshida et al., 1980; Yoshida et al., 1981; Cochrane et al., 1982). This year, the current leading anti-infection CSP-based vaccine against Pf malaria, RTS,S/AS01, has begun pilot implementation in Ghana, Malawi and Kenya. Notwithstanding, RTS,S/AS01 has shown to only provide rapidly waning protection in 50% of children and thus, intense research efforts are underway towards designing a more efficacious and durable anti-CSP vaccine (RTS,S Clinical Trials Partnership, 2015; Julien and Wardemann, 2019).

A molecular understanding of how the most potent monoclonal antibodies (mAbs) recognize sites of vulnerability on the parasite can guide next-generation vaccine design. PfCSP is composed of an N-terminal domain (N-CSP), a central repeat region comprising NANP motifs of varied number that are interspersed with related NVDP motifs, and a C-terminal domain (C-CSP) that comprises a linker region preceding an α-thrombospondin type-1 repeat (αTSR) domain (Fig. 1A). PfCSP is linked to the parasite membrane through a glycosylphosphatidylinositol anchor site. Numerous studies have shown that mAbs specific for the NANP repeat region and the junction immediately following N-CSP, which contains NANP motifs, NVDV motifs and the only copy of an NPDP motif, can mediate protection in animal models (Potocnjak et al., 1980; Yoshida et al., 1980; Foquet et al., 2014; Oyen et al., 2017; Triller et al., 2017; Kisalu et al., 2018; Tan et al., 2018; Imkeller et al., 2018; Murugan et al., 2019). The few described mAbs to C-CSP were functionally ineffective, probably, due to low accessibility of this domain on the sporozoite surface (Scally et al., 2018). In contrast, the functional relevance of N-CSP mAbs remains elusive. To date, no human mAb specific for this domain and only a handful of murine mAbs from immunization studies with recombinant Pf N-CSP have been reported (Espinosa et al., 2015; Herrera et al., 2015). These mAbs recognized N-CSP epitopes adjacent to Region I (RI; Fig. 1A), a site with high conservation across *Plasmodium* species, suggesting that RI may be a good target for cross-species vaccine development (Dame et al, 1984; Espinosa et al., 2015). Additionally, proteolytic cleavage of RI was linked to efficient sporozoite invasion of host hepatocytes (Espinosa et al., 2015; Coppi et al., 2005). Based on these observations, it has been proposed that adding N-CSP, including the RI motif, into a PfCSP subunit vaccine may improve protective efficacy compared to the current leading vaccine RTS,S/AS01, which lacks this domain. However, passive transfer of the most potent RI-targeting mAb 5D5 protected mice from infection in only one of the two tested transgenic rodent *P. berghei* (Pb) models that expressed a chimeric PbCSP with the Pf N-CSP domain (Espinosa et al., 2015), and its impact on Pf has not been determined. Thus, crucial information on how mAb 5D5 binds and inhibits Pf sporozoites is still missing.

**Figure 1.**
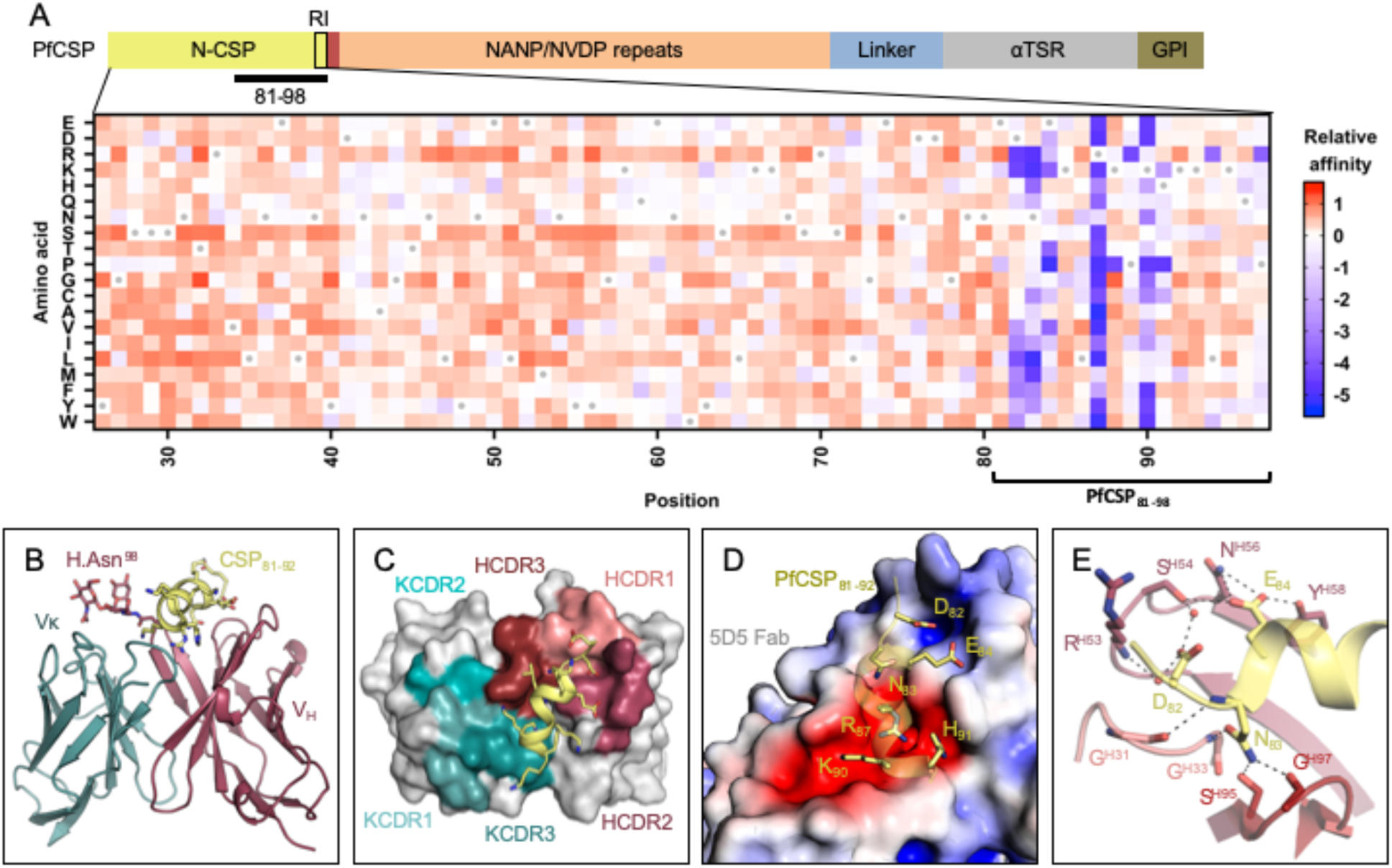
Molecular delineation of the mAb 5D5 epitope in PfCSP. (A, above) Schematic depicting the protein domain organization of PfCSP shown with the approximate location of Region I (RI) indicated by the black box and the junctional epitope represented by a dark red band. An approximate representation of PfCSP_81-98_ is illustrated by the black bar (not shown to scale). (Below) Heatmap of mAb 5D5 binding affinity for N-CSP single point mutant library. N-CSP residues included in PfCSP_81-98_ are indicated by the bracket at the bottom. The relative binding affinity is indicated by a diverging colour scale from red to blue, where red indicates a similar affinity while blue indicates decreased affinity. The X-axis denotes the N-CSP residue position and the Y-axis specifies the introduced single point mutations. Residues corresponding to the WT sequence are indicated by the gray dots. (B) Crystal structure showing the 5D5 Fab variable regions (heavy chain shown in red and kappa light chain shown in blue) bound to PfCSP N-terminal residues 81-92 (yellow), which are recognized in an α-helical conformation. The N-linked glycan on H.Asn98 of 5D5 Fab is represented as sticks. (C) mAb 5D5 CDRs contacting PfCSP. HCDRs 1, 2, and 3 (salmon, raspberry and firebrick red, respectively), and KCDRs 1 and 3 (light teal and deep teal, respectively) contribute to 5D5 Fab recognition, whereas KCDR2 (teal) does not. (D) Electrostatic surface potential of mAb 5D5 bound to PfCSP_81-98_. mAb 5D5 displays extensive shape and charge complementarity to PfCSP. Electrostatic calculations were performed using APBS (Baker et al., 2001) and rendered in Pymol (The PyMOL Molecular Graphics System, Version 2.0 Schrödinger, LLC); scale: -5 kT/e (red) to +5 kT/e (blue). (E) H-bonds (shown as black dashed lines) formed between mAb 5D5 HCDR residues and negatively charged PfCSP residues. Water molecules are shown as red spheres.

To gain a molecular understanding of how the mAb 5D5 recognizes PfCSP and inhibits Pf sporozoite infectivity, we solved the crystal structure of the 5D5 Fab in complex with a peptide derived from N-CSP and conducted in-depth binding and functional experiments with Pf sporozoites. We specifically quantified reactivity of mAb 5D5 to single live Pf sporozoites isolated from the midgut and salivary glands of mosquitoes using imaging flow cytometry, and tested its inhibitory potency against Pf sporozoites through *in vitro* traversal assays and *in vivo* passive mAb transfer experiments in mosquitoes. Our work provides a detailed molecular and functional understanding of mAb 5D5 recognition of its epitope, and highlights poor recognition of this N-CSP epitope on the Pf sporozoite surface.

## Results and Discussion

### mAb 5D5 binds an α-helical motif in N-CSP

To understand the molecular basis for mAb 5D5 recognition of PfCSP, we solved the crystal structure of the 5D5 Fab in complex with PfCSP_81-98_ to 1.85 Å resolution (Supplementary Table 1). We specifically selected PfCSP residues 81-98 for our studies to ensure inclusion of the mAb 5D5 epitope, identified as Pf N-CSP residues 82-91 by yeast display epitope mapping (Fig. 1A and S1A) in agreement with previous reports (Espinosa et al., 2015), as well as conserved RI residues KLKQP in positions 93-97. Consistent with our yeast display experiments, we observed strong electron density for N-CSP residues 81-92 (EDNEKLRKPKHK) in the crystal structure (Fig. S1B). PfCSP residues 83-91 formed an α-helix when bound by 5D5 Fab (Fig. 1B), in line with secondary structure predictions based on the primary sequence (Drozdetskiy et al., 2015). Importantly, while structures of a variety of polypeptides derived from PfCSP (the junctional region following N-CSP, the NANP repeat region, and the C-terminal αTSR domain) have been solved in complex with a broad range of antibodies (Oyen et al., 2017; Imkeller et al., 2018; Scally et al., 2018; Tan et al., 2018; Kisalu et al., 2018; Julien and Wardemann, 2019; Murugan et al., 2019), our crystal structure of the 5D5 Fab in complex with PfCSP_81-98_ provides the first insight into the subdomain architecture of Pf N-CSP. However, further studies are needed to elucidate the conformation of residues comprising RI, which were disordered and unresolved in our crystal structure, as well as the overall structure of Pf N-CSP.

mAb 5D5 contacts PfCSP with all complementarity-determining regions (CDRs) except kappa light chain CDR 2 (KCDR2; Fig. 1C). The heavy chain CDRs (HCDRs) form the majority of interactions with 498 Å^2^ buried surface area (BSA) compared to 160 Å^2^ BSA for the kappa light chain. Furthermore, the mAb 5D5 CDRs possess extensive shape and electrostatic complementarity to this highly charged N-CSP epitope (Fig. 1D). An electropositive pocket formed by HCDR2 contacts PfCSP residues D82 and E84 via H-bonds with Ser54^Oγ^, Asn56^Nδ2^, and Tyr58^OH^, and water-mediated H-bonds with Arg53^N^ and Ser54^Oγ^ (Fig. 1E). Additionally, an electronegative pocket formed by HCDR2, HCDR3, KCDR1, and KCDR3 contacts PfCSP electropositive residues R87, K90 and H91 via several H-bonds and salt bridges (Fig. 2A). The significance of this shape and charge complementarity for high affinity binding was observed in our yeast display experiments, as mutations maintaining both side chain length and electrostatic properties, such as K90R, were more likely to sustain high affinity binding than those that did not (K90E, K90D; Fig. 1A).

**Figure 2.**
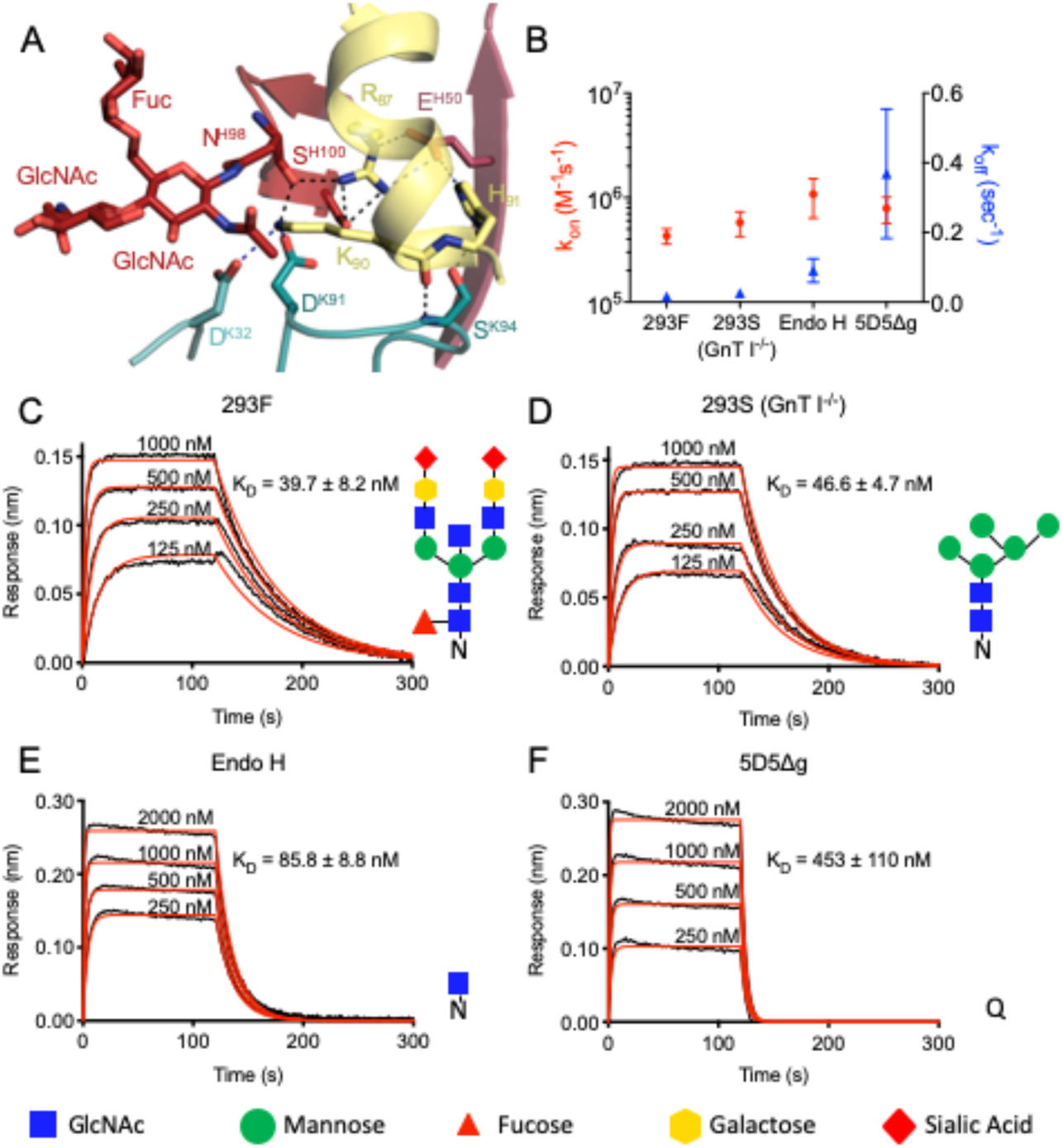
5D5 paratope glycosylation mediates high affinity binding. (A) Interactions formed by mAb 5D5 H.Asn98-linked GlcNAc moiety and surrounding CDR residues with PfCSP. H-bonds are shown as black dashed lines, and salt bridges are shown as blue dashed lines. (B-F) Binding kinetics of twofold dilutions of (C) 293F, (D) 293S (GnT I^-/-^), (E) Endo H and (F) 5D5Δg Fab glycoform variants to full length PfCSP. (B) Mean k_on_ and k_off_ rates of the 5D5 Fab glycoform variants binding to full length PfCSP are plotted on the left and right y-axis, respectively. Mean k_on_ rates are shown as red circles, and mean k_off_ rates as blue triangles. (C-F) Representative sensorgrams are shown in black and 1:1 model best fits in red. Mean K_D_ values are as listed. K_D_ values, and k_on_ and k_off_ rates were determined by FortéBio’s Data Analysis software 9.0. Standard error values are reported as the standard deviation. Data are representative of three independent measurements. Corresponding glycan structures are shown using symbols adhering to the Symbol Nomenclature for Glycans (Varki et al., 2015).

Consistent with prediction from the primary mAb 5D5 sequence, we observed electron density for two GlcNAc and one α1-6Fuc moieties indicative of N-linked glycosylation at Asn98 of HCDR3 (Fig. 2A and S1C). Importantly, the first N-linked GlcNAc moiety contacts the aliphatic portion of K90 of PfCSP_81-98_, conferring 48 Å^2^ of BSA, while the other sugars did not interact with the peptide. In this way, the paratope glycan contributes to mAb 5D5 occlusion of PfCSP residue K90. Interestingly, K90 is one of four lysine residues (including K85, K88, and K92) directly upstream of RI that have previously been proposed to be important for binding heparan sulfate proteoglycans on the surface of hepatocytes to initialize liver invasion (Zhao et al., 2016). Notably, the α-helical conformation adopted by N-CSP residues 83-91 upon mAb 5D5 binding positions the remaining three lysine residues, K85, K88 and K92, on the same exposed face of the helix (Fig. S1B). Thus, our molecular description of PfCSP recognition by mAb 5D5 demonstrates optimal antibody-antigen characteristics associated with high affinity binding to a putative functional site on Pf sporozoites.

### mAb 5D5 paratope glycosylation is critical for high affinity recognition of recombinant PfCSP

To determine whether the N-linked glycan on H.Asn98 affects mAb 5D5 binding, we generated four different forms of the mAb 5D5 glycan and measured their binding kinetics to full-length PfCSP using biolayer interferometry (BLI; Fig. 2B-F). Specifically, we generated four 5D5 Fab variants with either: 1) a complex glycan, as in the crystal structure (by expression in HEK 293F cells; 293F); 2) a high mannose glycan (by expression in HEK 293S (GnT I^-/-^) cells; 293S (GnT I^-/-^)); 3) a single GlcNAc moiety (by expression in HEK 293S (GnT I^-/-^) cells followed by Endo H treatment; Endo H); or 4) an H.N98Q mutation removing the N-linked glycosylation site altogether (5D5Δg). The 293F, 293S (GnT I^-/-^) and Endo H-treated 5D5 Fabs bound with high affinity to full-length PfCSP with KD’s of 39.7 ± 8.2 nM, 46.6 ± 4.7 nM and 85.8 ± 8.8 nM, respectively (Fig. 2C-E). In contrast, the 5D5Δg mutant Fab bound PfCSP with weaker affinity (KD of 453 ± 110 nM) due to an 11-fold faster off-rate compared to HEK 293F-expressed 5D5 Fab (Fig. 2B and F), while the on-rates of all glycoform Fabs remained within the same order of magnitude. Together with the crystal structure, these results underline the importance of the H.Asn98-linked GlcNAc moiety for high affinity mAb 5D5 binding to recombinant PfCSP, and highlight a rare occurrence for such a post-translational modification to participate in the antibody-antigen interaction and improve the kinetics of antigen binding.

### mAb 5D5 does not efficiently bind or inhibit salivary gland Pf sporozoites

The role of the N-linked glycan on H.Asn98 in the binding of 5D5 IgG to freshly isolated salivary gland Pf sporozoites was quantified by imaging flow cytometry. As these preparations contained both live and dead sporozoites, we focused our analyses on live sporozoites that were negative for propidium iodide staining (Fig. S2). As positive and negative controls, we used human mAbs targeting the PfCSP central repeat (1210; Imkeller et al., 2018) and C-CSP (1710; Scally et al., 2018), respectively. In line with previous reports, mAb 1710 failed to recognize mature Pf sporozoites isolated from mosquito salivary glands, whereas mAb 1210 strongly bound these sporozoites (Fig. 3A-D). We detected a two-fold decrease in mean fluorescence intensity (MFI) between mAbs 1210- and 5D5- or 5D5Δg-bound sporozoites (Fig. 3C). The observed differences can be explained by the frequencies of the targeted epitopes on the sporozoite surface. Indeed, mAb 1210 likely binds repeated NANP motifs within the central region whereas mAbs 5D5 and 5D5Δg can only react with a single N-CSP motif. In contrast to previous reports, we found that mAbs 5D5 did not bind the majority of sporozoites (Fig. 3B and D). Mutation of the glycosylation site further decreased the proportion of 5D5Δg-bound sporozoites from 27 to 13% (Fig. 3D). These results demonstrate the importance of mAb 5D5 paratope glycosylation for PfCSP binding on the sporozoite surface. However, they also reveal low levels of overall reactivity of this antibody to live salivary gland Pf sporozoites.

**Figure 3.**
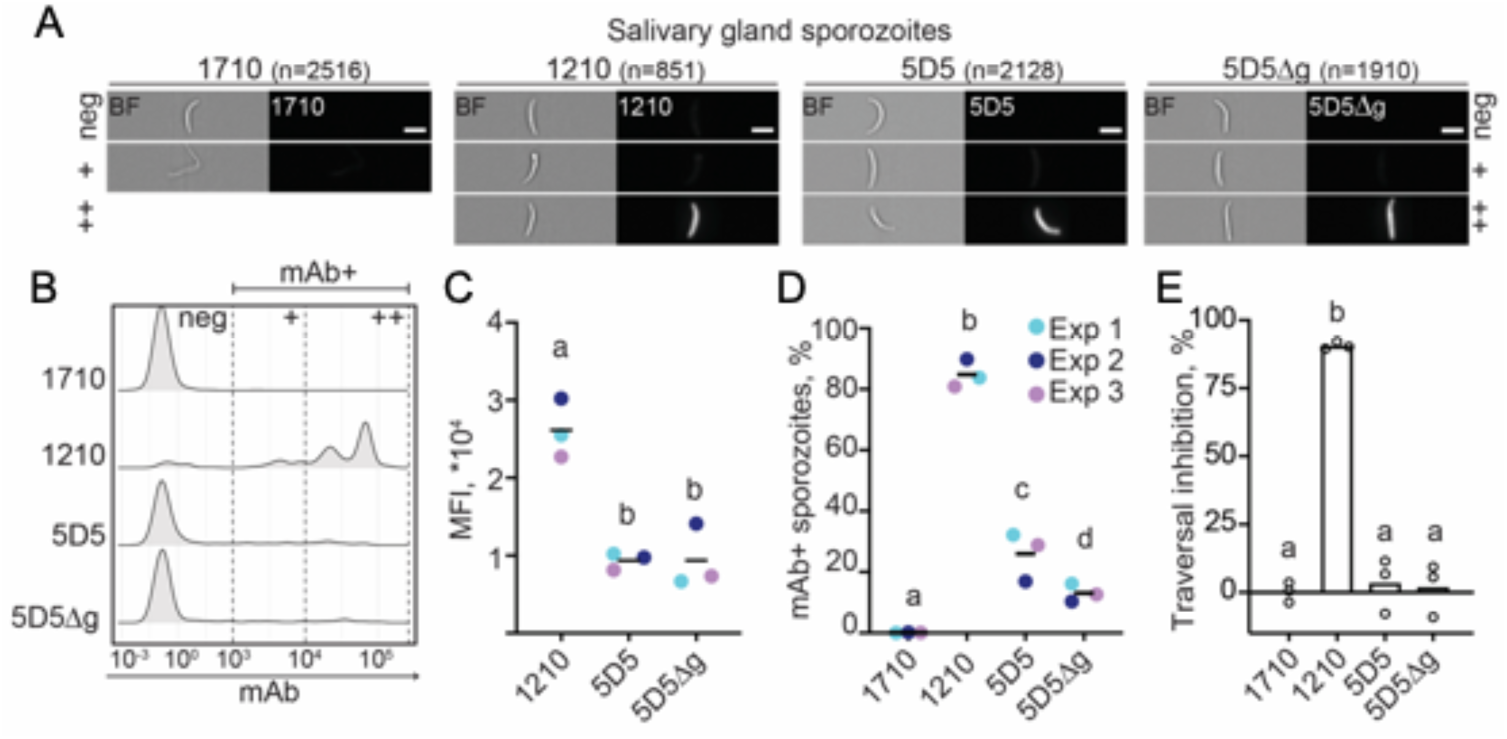
mAb 5D5 binding to salivary gland sporozoites and traversal inhibition activity. (A-D) Imaging flow cytometry of live salivary gland sporozoites isolated from mosquito thorax after incubation with human mAb 1710 or 1210 (negative and positive controls, respectively), or mAb 5D5 or 5D5Δg. (A) Representative images of sporozoites in brightfield (BF, left panels) and mAb-bound fluorescent sporozoites (right panels). Scale bars - 5 µm. Total number of sporozoites analysed per condition is indicated in parentheses (N=3). (B) Comparative density plots of a representative experiment showing the fluorescence intensities of three arbitrarily-designated groups of mAb-bound sporozoites (neg – negative, + – low intensity, ++ – high intensity). (C) Mean fluorescence intensities (MFI) of the mAb-positive sporozoites. (D) Quantification of mAb-positive sporozoites (%). (C-D) Colors show results of three independent experiments. (E) Results of mAb inhibition of sporozoites in *in vitro* traversal assay tested at 100 µg/mL mAb concentration (N=3). Statistically significant differences (p<0.05) between the groups are indicated by different letters (z-test (C and D); paired Friedman test followed by Dunn’s *post-hoc* test (E)).

We next evaluated how the low sporozoite binding observed for mAb 5D5 translated into inhibitory potency against Pf sporozoites in a hepatocyte traversal assay. In line with the mAb binding efficiencies, only mAb 1210 completely blocked sporozoite traversal of hepatocytes, whereas mAb 5D5 was as inefficient at inhibiting traversal as negative control mAb 1710, regardless of the presence of the paratope glycan (Fig. 3E). Poor mAb 5D5 binding to sporozoites and lack of traversal inhibition precluded further functional testing of mAb 5D5 in Pf infections in a sophisticated humanized mouse model. We conclude that, in spite of high affinity interaction with the N-CSP epitope in recombinant PfCSP, the overall low levels of mAb 5D5 binding to live Pf sporozoites preclude efficient inhibition of parasite traversal.

### mAb 5D5 does not inhibit *in vivo* sporozoite development in mosquitoes

As CSP is essential for sporozoite development in mosquitoes (Menard et al., 1997), we extended our antibody binding and functional examination to immature Pf sporozoites isolated from oocysts in the mosquito midgut (Fig. 4A). Similar to our observations with mature sporozoites, mAb 5D5 exhibited low binding efficiency to immature sporozoites, as measured by MFI and percentage of mAb-positive sporozoites determined using imaging flow cytometry (Fig. 4B-C). Also consistent with our findings with salivary gland sporozoites, paratope glycosylation increased the proportion of mAb 5D5-bound midgut sporozoites by two-fold (Fig. 4D).

**Figure 4.**
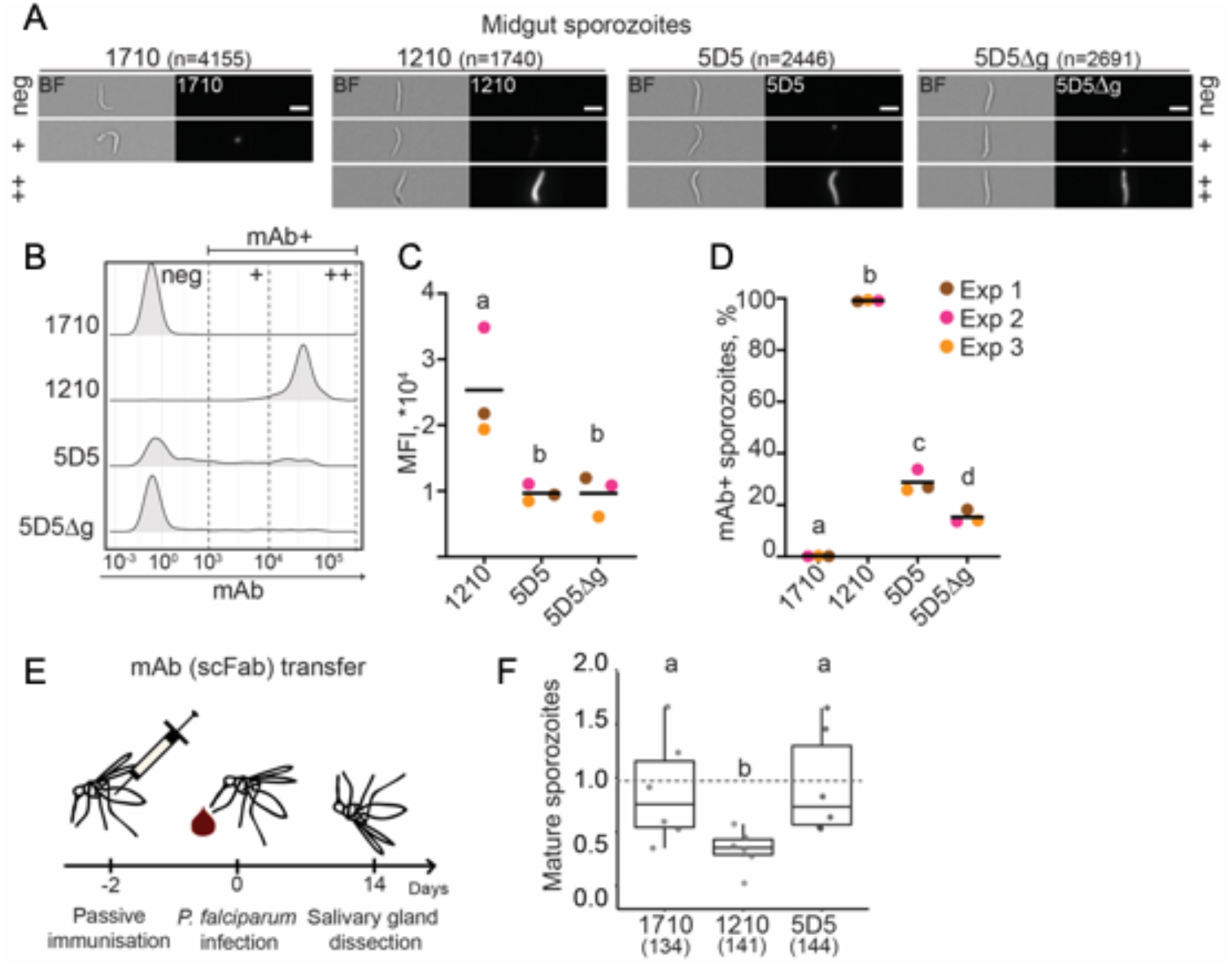
mAb 5D5 binding to midgut sporozoites and inhibition of sporogonic development within mosquitoes. (A-D) Imaging flow cytometry of live midgut sporozoites isolated from oocysts after incubation with human mAb 1710 or 1210 (negative and positive controls, respectively), or mAb 5D5 or 5D5Δg. (A) Representative images of sporozoites in brightfield (BF, left panels) and mAb-bound fluorescent sporozoites (right panels). Scale bars - 5 µm. Total number of sporozoites analysed per condition is indicated in parentheses (N=3). (B) Comparative density plots of a representative experiment showing the fluorescence intensities of three arbitrarily-designated groups of mAb-bound sporozoites (neg – negative, + – low intensity, ++ – high intensity). (C) Mean fluorescence intensities (MFI) of the mAb-positive sporozoites. (D) Quantification of mAb-positive sporozoites (%). (C-D) Colors show results of three independent experiments. (E) Schematic representation of passive single-chain Fab (scFab) transfer by mosquito injection. (F) Results of scFab transfer experiments expressed relative to control PBS-injected mosquitoes (N=6, total mosquito numbers analyzed are indicated below in parentheses). The box plots show the upper and lower quantiles and the median of the distribution. Each dot represents normalized sporozoite loads in one experiment. Statistically significant differences (p<0.05) between the groups are indicated by different letters (z-test (C and D); maximum likelihood estimation (MLE)(F)).

To evaluate the inhibitory activity of mAb 5D5 against Pf sporozoites in their natural environment *in vivo*, we developed a passive mAb transfer assay for mosquitoes that examined sporozoite maturation and salivary gland invasion. Mosquitoes were injected with recombinant single-chain Fabs (scFabs) two days before Pf infection, and sporozoite loads in dissected salivary glands were quantified two weeks later (Fig. 4E). Injection of scFab1210 significantly reduced the number of mature sporozoites in the salivary glands. In contrast, transfer of scFab1710 or scFab5D5 did not affect sporozoite development and invasion (Fig. 4F). Taken together, these results demonstrate that mAb 5D5 fails to efficiently recognize its epitope on the surface of Pf parasites, and lacks inhibitory potency against sporozoites in both the vector and the host.

### Concluding Remarks

In this report, we demonstrate that despite high-affinity binding to recombinant PfCSP, mAb 5D5 does not recognize the majority of live Pf sporozoites, indicating that its epitope is not readily accessible or present on the sporozoite surface. Consequently, as shown in the current study, mAb 5D5 is unable to block Pf sporozoite development in the mosquito or inhibit sporozoite traversal of hepatocytes. The lack of potent human N-CSP-specific mAbs in multiple screens based on full-length recombinant PfCSP baits or unbiased antigen-agnostic approaches (Fig. S3; Triller et al., 2017; Murugan et al., 2018; Tan et al., 2018; Kisalu et al., 2018; Julien and Wardemann, 2019) is in line with these observations. Overall, to date, there is little evidence to support N-CSP as a source of potent or protective epitopes to block Pf infection. Therefore, the repeating motifs in the central domain and N-terminal junction remain the most promising PfCSP regions for anti-infective vaccine design to elicit protective mAbs.

## Materials and Methods

### Mutant N-CSP yeast display library construction and transformation

Epitope mapping using phage display was adapted from a previously published method (Van Blarcom et al., 2015). Construction of the mutant library required generation of a linearized vector and a library of mutant N-CSP inserts. The mutant insert library was generated by two rounds of PCR using primers that carried the randomized codon “NNK” (Tables S2 and S3), and mixing the products (Amplicons 1 to 9) at an equal molar ratio. The vector was generated by overlapping PCR using vector-F/R primers (Table S2). All PCR products were amplified using KOD DNA polymerase (EMD Millipore) and purified by gel extraction (Clontech Libraries).

The EBY100 yeast strain was purchased from ATCC. The yeast vector was generated by modification of the commercially available pCTcon2 vector (Addgene; Chao et al., 2006). The mutant N-CSP insert was cloned with N-terminal V5 and C-terminal HA epitope tags. The Aga2p yeast protein gene was inserted downstream of the HA epitope tag to allow for yeast surface display of the N-CSP (plasmid pCTcon2-rsCSP-V5-HA-Aga2p). Yeast transformation was performed as described previously (Benatuil et al., 2010). In summary, 4 μg of the linearized yeast expression vector and 8 μg of the N-CSP mutant library insert were used for transformation. Transformants were plated on SDCAA plates and incubated at 30°C for 2 days. Over 10^8^ colonies were collected, resuspended in YPD media with 15% glycerol, and stored at −80°C until use.

### 5D5 Fab production and purification

mAb 5D5 V_L_ and V_H_ regions were individually cloned into pcDNA3.4-TOPO expression vectors immediately upstream of human Igκ and Igγ1-CH1 domains, respectively. The resulting 5D5 Fab light and heavy chain vectors were co-transfected into either HEK293F or HEK293S (GnT I^-/-^) cells for transient expression, and purified via KappaSelect affinity chromatography (GE Healthcare), cation exchange chromatography (MonoS, GE Healthcare), and size exclusion chromatography (Superdex 200 Increase 10/300 GL, GE Healthcare). For binding studies, 5D5 Fab expressed in HEK293S (GnT I^-/-^) cells was digested with Endoglycosidase H, followed by an additional size exclusion chromatography step (Superdex 200 Increase 10/300 GL, GE Healthcare). Lastly, the 5D5Δg Fab was produced by site-directed mutagenesis of the mAb 5D5 V_H_ region using Accuprime Pfx Supermix (Thermo Fisher Scientific). 5D5Δg Fab was expressed in HEK293F cells and purified by chromatography as described above.

### IgG production and purification

For yeast display experiments, mAb 5D5 was produced in ExpiCHO cells as a mouse IgG1 with AVI tag for biotinylation. The IgG was then purified using protein G affinity chromatography (HiTrap Protein G HP, GE Healthcare) and size-exclusion chromatography (Superdex 200, GE Healthcare). Biotinylation was performed as previously described (Ekiert et al., 2011).

For production of 5D5 and 5D5Δg IgGs for non-yeast display experiments, site-directed mutagenesis was performed using In-Fusion (Takara Bio) on the pcDNA3.4-TOPO vectors encoding the 5D5 Fab heavy chain and 5D5Δg Fab heavy chain sequences to substitute two stop codons with two residues (DK), allowing for expression of the Igγ1-CH2 and Igγ1-CH3 domains. 5D5 IgG, 5D5Δg IgG, 1710 IgG (Scally et al., 2018), 1210 IgG (Imkeller et al., 2018), and IgGs elicited by the PfSPZ-CVac Challenge (Mordmüller et al., 2017; Murugan et al., 2018) were transiently expressed in HEK293F cells by co-transfection of paired Ig heavy and light chains, and purified through protein A affinity chromatography (GE Healthcare), followed by size exclusion chromatography (Superdex 200 Increase 10/300 GL, GE Healthcare).

### scFab production and purification

scFab constructs were designed by cloning paired light and heavy chains, separated by a 72-residue linker, into a pcDNA3.4-TOPO expression vector. The resulting constructs were transiently expressed in HEK293F cells, and purified by KappaSelect affinity chromatography (GE Healthcare), followed by size exclusion chromatography (Superdex 200 Increase 10/300 GL, GE Healthcare).

### Recombinant PfCSP production and purification

A construct of full-length PfCSP isolated from strain NF54 (UniProt accession no. P19597, residues 20-384) was designed with potential N-linked glycosylation sites mutated to glutamine (Scally et al., 2018). The resulting construct was transiently transfected in HEK293F cells, and purified by HisTrap FF affinity chromatography (GE Healthcare) and size exclusion chromatography (Superdex 200 Increase 10/300 GL, GE Healthcare).

A construct encoding PfCSP residues 71-104 was cloned into a pETM-22 vector. Competent BL21(DE3) *E. coli* cells were transformed with the resulting plasmid and cultured to an optical density of approximately 0.6-0.8. Expression of PfCSP_71-104_ was induced using 1 mM isopropyl β-D-1-thiogalactopyranoside (IPTG). Approximately 4 h after induction, cells were lysed by sonication, and purified through HisTrap affinity chromatography (GE Healthcare) and size exclusion chromatography (Superdex 75 10/300 GL, GE Healthcare).

### Yeast display epitope mapping

For each sorting round, ∼10^9^ yeast cells from the frozen stock were cultured in 250 mL of SDCAA media for 16 h at 27.5°C until an OD of 1.9 was reached. Cells were pelleted, resuspended in 35 mL of SGR-CAA induction media and incubated for 30 h at 18°C until an OD of 1.4 was reached. After harvesting approximately 8 mL of cell culture, the pellet was washed 3 times with PBS and finally resuspended in 5 mL of PBS. Biotinylated 5D5 IgG was incubated with BB515-streptavidin at a molar ratio of 1:2 for 20 min. The biotinylated 5D5 IgG-streptavidin BB515 complex and Anti-HA PE antibody were added to the 5 mL of resuspended yeast cells with a final concentration of 20 nM for each stain, followed by overnight incubation at 4°C with head-to-head rotation in the dark. Next, cells were washed twice with PBS, resuspended in 5 mL of PBS, and sorted at the TSRI Flow Cytometry Core Facility. Two gates were applied for simultaneous sorting (Fig. S1A): one where binding of 5D5 IgG was completely abrogated (PE only) and one where binding was unaffected (PE and BB515). The second round of sorting saw significant enrichment in either gate.

### Deep mutational scanning data analysis

Sequencing data were obtained in FASTQ format and parsed using SeqIO module in BioPython (Cock et al., 2009). After trimming the primers, a paired-end read was then filtered and removed if the corresponding forward and reverse reads were not reverse-complemented. The position of the randomized codon was then identified by the internal barcode. Each mutation was called by comparing individual paired-end reads to the wild type (WT) reference sequence. Frequency of mutation *m* in sample *s* was computed as:

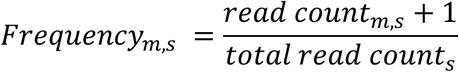

Relative affinity of mutation *m* was computed as:

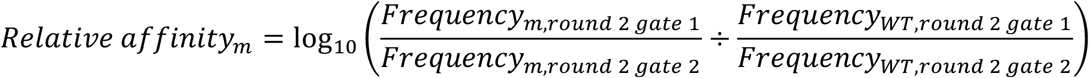

The pseudo read count of 1 in the calculation of frequency was to prevent division by zero during the calculation of relative affinity. Relative affinity of WT is 0.

Raw sequencing data have been submitted to the NIH Short Read Archive under accession number: BioProject PRJNA578947. Custom python scripts for analyzing the deep mutational scanning data have been deposited to https://github.com/wchnicholas/CSP_Nterm_yeast_display.

### Crystallization and structure determination

Purified 5D5 Fab and CSP 81-98 peptide (GenScript) were mixed in a 1:5 molar ratio. The 5D5 Fab/CSP 81-98 complex was then mixed in a 1:1 ratio with 20% (w/v) PEG 3350, 0.2 M di-ammonium citrate. Crystals appeared after ∼20 h, and were cryoprotected in 15% (v/v) ethylene glycol before being flash-frozen in liquid nitrogen. Data were collected at the 08ID-1 beamline at the Canadian Light Source, processed and scaled using XDS (Kabsch, 2010). The structure was determined by molecular replacement using Phaser (McCoy et al., 2007) and a Fab model from our internal database as the search model. Refinement of the structure was performed using phenix.refine (Adams et al., 2010) and iterations of refinement using Coot (Emsley et al., 2010). The crystal structure has been deposited in the Protein Data Bank (PDB ID 6UUD).

### BLI binding studies

BLI (Octet RED96, FortéBio) experiments were conducted to determine the binding kinetics of the 5D5 Fab glycoform variants to recombinant PfCSP diluted to 10 µg/mL in kinetics buffer (PBS, pH 7.4, 0.01% [w/v] BSA, 0.002% [v/v] Tween-20) was immobilized onto Ni-NTA (NTA) biosensors (FortéBio). After a steady baseline was established, biosensors were dipped into wells containing twofold dilutions of each 5D5 Fab glycoform variant in kinetics buffer. Tips were then immersed back into kinetics buffer for measurement of the dissociation rate. Kinetics data were analyzed using FortéBio’s Data Analysis software 9.0, and curves were fitted to a 1:1 binding model. Mean kinetic constants and corresponding standard deviation values are reported as the result of three independent experiments for each 5D5 Fab glycoform variant.

BLI experiments were also done to determine the avidity of IgGs isolated from the PfSPZ-CVac Challenge (Mordmüller et al., 2017) to full length recombinant PfCSP and N-CSP construct, PfCSP_71-104_. Unrelated malaria protein Pfs25 was used to block non-specific binding and 5D5 IgG was used as a positive control. PfCSP_71-104_, PfCSP or Pfs25 was diluted to 10 µg/mL in kinetics buffer and immobilized onto Ni-NTA (NTA) biosensors (FortéBio). Once a stable baseline was established, biosensors were dipped into wells, each containing a different IgG diluted to 500 nM in kinetics buffer. Tips were subsequently dipped back into kinetics buffer to observe any dissociation of IgG.

### Mosquito rearing, parasite infections and sporozoite isolations

*Anopheles coluzzii* (Ngousso strain) mosquitoes were maintained at 29°C 70**–**80% humidity 12/12 h day/night cycle. For *P. falciparum* infections, mosquitoes were fed for 15 min on a membrane feeder with NF54 gametocytes cultured with O^+^ human red blood cells (Haema, Berlin), and, thereafter, kept in a secured S3 laboratory according to the national regulations (Landesamt für Gesundheit und Soziales, project number 297/13). The *P. falciparum* NF54 clone used in this study originated from Prof. Sauerwein’s laboratory (RUMC, Nijmegen) and was authenticated for *Pfs47* genotype by PCR on genomic DNA. *P. falciparum* asexual cultures were monthly tested for *Mycoplasma* contamination. Unfed mosquitoes were removed shortly after infections. Blood fed mosquitoes were offered an additional uninfected blood meal eight days post infection, maintained at 26°C for 12 and 14/15 days, and used for the midgut and salivary gland dissections, respectively. The midgut or salivary gland sporozoites were isolated into HC-04 complete culture medium (MEM without L-glutamine (Gibco) supplemented with F-12 Nutrient Mix with L-glutamine (Gibco) in 1:1 ratio, 15 mM HEPES, 1.5 g/L NaHCO3, 2.5 mM additional L-glutamine, 10% FCS) and kept on ice until further use.

### Imaging flow cytometry of sporozoites

Isolated sporozoites were diluted in PBS/1% FCS to 3×10^6^/mL and incubated for 30 min at 4°C with 1 µg/mL recombinant IgGs, washed (16,000 × g, 4 min, 4°C) and incubated with Cy5-conjugated anti-human IgG1 (0.4 µg/ml, DRFZ Core Facility, Berlin) for 30 min at 4°C. After incubation with the secondary antibody, sporozoites were further incubated with propidium iodide (20 µg/mL, Sigma Aldrich) for 5 min at room temperature as previously described (Costa et al., 2018). Sporozoites were acquired after one wash in PBS using the ImageStreamX Mk II instrument (Merck Millipore) with a 60X objective for 15-20 min per sample. The experiments were performed in three replicates. To avoid a possible bias due to the variable pre-acquisition waiting times on ice, the order of samples was swapped between the experimental replicates. Quantification of propidium iodide staining was performed using the Intensity_MC_Ch04, whereas mAb binding was quantified by Cy5-conjugated secondary antibody signal Intensity_MC_Ch11. Single sporozoites were manually selected by brightfield images (Channel 9) and only live, propidium iodide-negative sporozoites were gated for mAbs binding efficiency analysis (for gating strategy see Fig. S2). The analysis was performed with IDEAS 6.2 (Merck Millipore). Raw data were exported as .txt files and represented in dot and density plots using RStudio Version 1.1.453.

### Pf sporozoite hepatocyte traversal assay

Pf traversal assays were performed as previously described (Triller et al., 2017). In brief, the salivary gland sporozoites were isolated from mosquito thorax and treated with mAbs (100 µg/mL) for 30 min on ice. The sporozoite preparations were seeded on human hepatocytes (HC-04; Sattabongkot et al., 2006) for 2 h at 37°C and 5% CO2 in the presence of dextran-rhodamine (0.5 mg/mL) (Molecular Probes). mAb-untreated Pf sporozoites were used to measure the maximum traversal capacity. Cells incubated only with uninfected mosquito thoracic material were used as a background control. Cells were washed and fixed with 1% (v/v) formaldehyde in PBS. Dextran positivity was detected by FACS LSR II instrument (BD Biosciences). Data analysis was performed by background subtraction and normalization to the maximum traversal capacity of mAb-untreated Pf sporozoites using FlowJo v.10.0.8 (Tree Star).

### Passive transfer of scFabs into mosquitoes

1-2-day-old female *A. coluzzii* mosquitoes were injected on ice with 100 ng (285 µL) of scFab diluted in PBS or with 285 µL PBS as an injection control. Two days later, mosquitoes were infected with *P. falciparum* NF54 following the protocol described above. Mosquito heads were carefully pulled off 14 days later and the attached salivary glands were collected and washed with PBS. Dissected salivary glands were pooled for each group, homogenized and the freshly isolated sporozoites were counted using a Malassez hemocytometer. The average number of sporozoites per mosquito was calculated for each group.

### Enzyme-linked immunosorbent assay

Antigen ELISA detHigh binding 384 well polystyrene plates (Corning) were coated with recombinantly expressed PfCSP_71-104_ comprising N-CSP at 50 ng/well overnight at 4°C. 1% BSA in PBS was used for blocking the wells at RT. Binding of mAbs to N-CSP was determined by incubating the coated plates with serially diluted mAb at 4.00, 1.00, 0.25, 0.06 µg/mL concentrations. Bound mAb was detected using goat anti–human IgG-HRP (Jackson ImmunoResearch) at 1:1,000 dilution in 1% BSA in PBS and One-step ABTS substrate (Roche). Non-PfCSP reactive antibody, mGO53, was used as negative control (Wardemann et al., 2003). Area under the curve (AUC) based on diluted antibody series was calculated using GraphPad Prism 7.04 (GraphPad).

### Statistical analysis

No samples were excluded from the analyses. Mosquitoes from the same batches were randomly allocated to the experimental groups (age range: 1–2 days). The experimenters were not blinded to the group allocation during the experiment and/or when assessing the outcome. Sample sizes were chosen according to best practices in the field and previous experience (Costa et al., 2018).

For Figures 3C and 4C, Mean Fluorescence Intensity (MFI) and standard error of the mean (SEM) of the mAb-bound live sporozoites were first computed from the data. We then associated an MFI and standard error to each treatment by computing the average MFI across the three independent experiments and subsequently computing the standard deviation (STD) as STD = [(SEM_1_^2^ + SEM_2_^2^ + SEM_3_^2^)^1/2^]/3. The null hypothesis was that the MFIs of sporozoites bound by the tested mAbs were not different. Due to the large sample sizes examined, a z-test was used to compare the MFI of the three conditions, and the obtained p-values are summarized below with significant p-values highlighted in green:

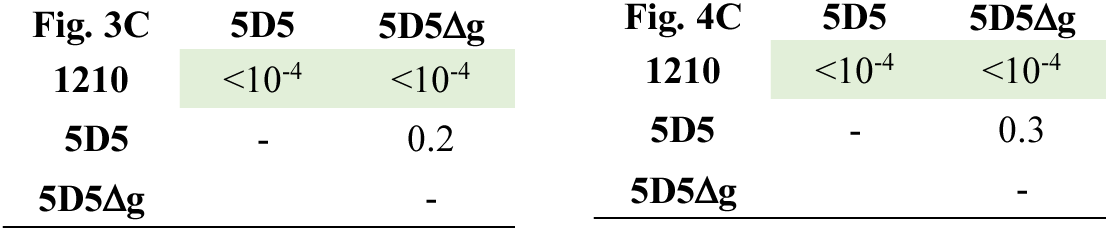

The combined *p*-values from the z-test were much smaller than the total sample size, which is roughly 10^4^. *p*-values much smaller than the inverse of the population size were therefore rounded up to 10^-4^.

For Figures 3D and 4D, sample sizes (*tot1*) and proportion of mAb-bound sporozoites per experiments (*pos1*) for each independent experiment (N=3) are summarized below:

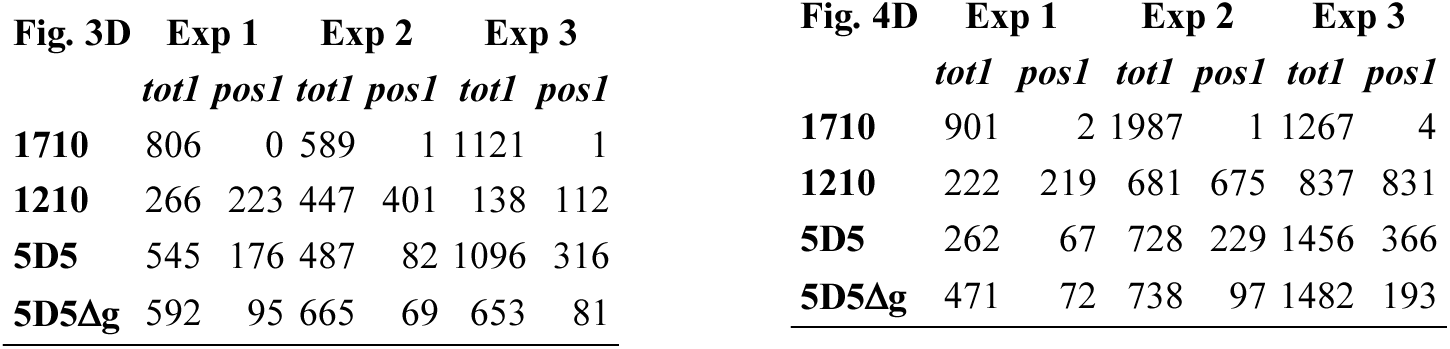

The null hypothesis was that the proportions of sporozoites bound by the tested mAbs were not different. Normality was verified and a z-test was used to compare the fractions of mAb-bound sporozoites. We first computed the fraction *f* of mAb-bound for each mosquito tissue, experiment and treatment as *f_i_* = pos_*i*_/tot_*i*_. The fraction *f* can be considered as the probability that a sporozoite taken at random is bound by a certain mAb. The error associated to *f* is therefore given by 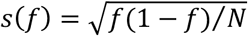, where N is the sample size given in the column *tot1*. Within each experiment, we used a two-sided z-test to test the null hypothesis that the fractions *f* associated to the tested mAbs were not different, resulting in six pairwise comparisons per experiment. The resulting *p*-values from the three independent experiments were combined using Fisher method and are summarized below with significant *p*-values highlighted in green:

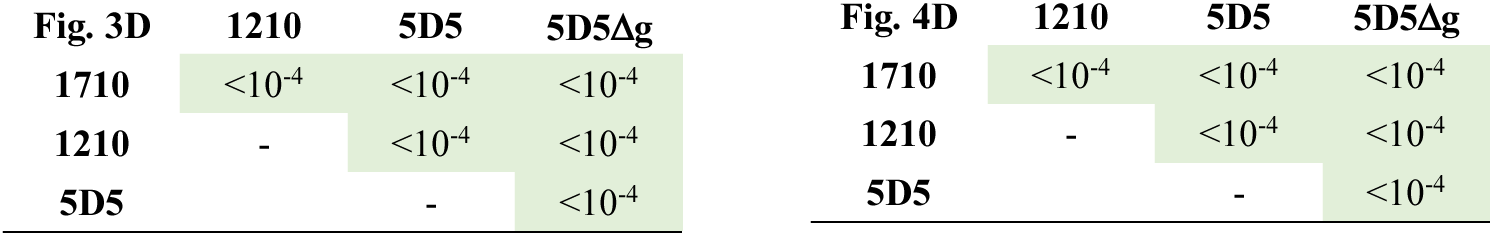

For both Figures 3D and 4D, the fractions *f* organized from strong to weak are as follows: 1210, 5D5, 5D5Δg, 1710. The combined *p*-values computed with the Fisher method are much smaller than the total sample size, which is roughly 10^4^. The reported *p*-values are therefore the inverse of the total sample size.

Statistical analysis in Figure 3E was performed using GraphPad Prism 8 (paired Friedman test followed by Dunn’s post-hoc test, paired values for mAb treatment per experiment) and p-values below 0.05 were considered significant (*p < 0.05).

For Figure 4F, number of dissected mosquitoes and mean number of sporozoites per mosquito in each independent experiment (N=6) are summarized below:

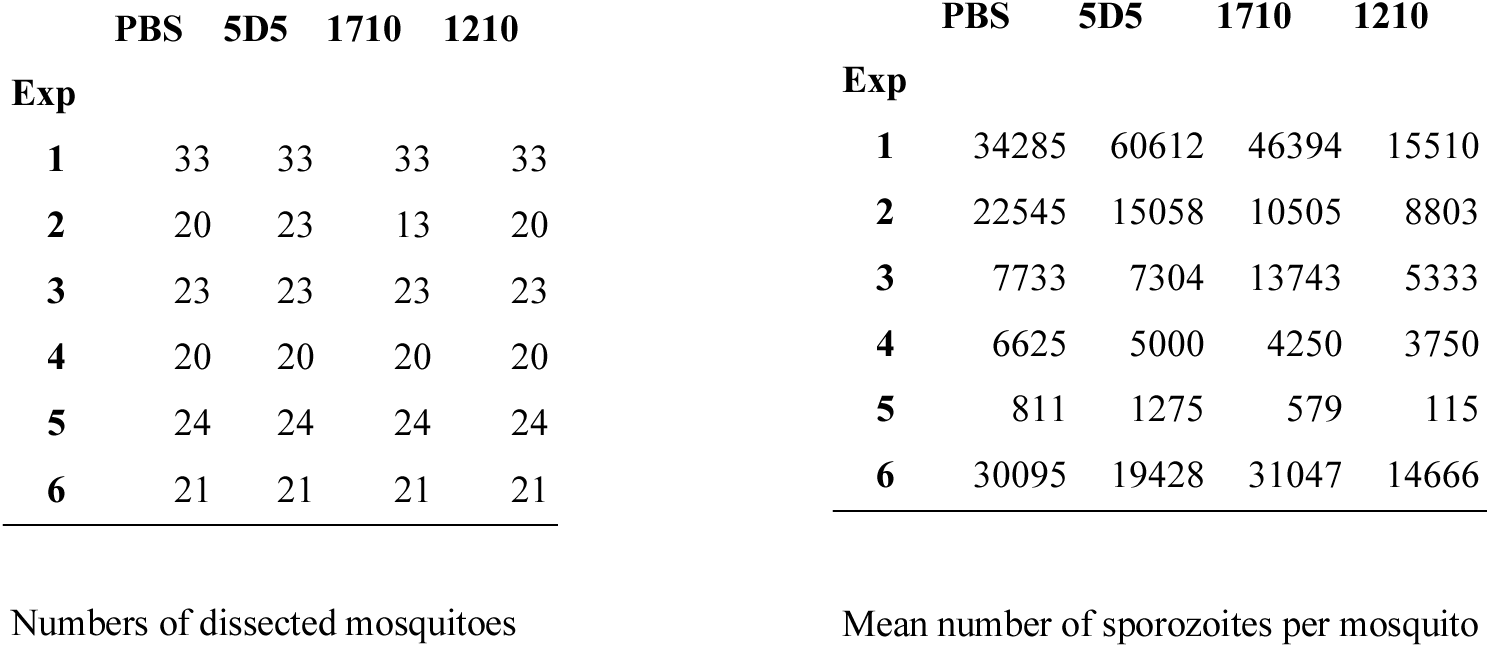

The null hypothesis was that the average number of sporozoites for each scFab and for each experiment independently was not significantly different. To perform this test, we used the number of oocysts per mosquito from the same experiments (data available upon request). We have assumed that the number of oocysts per mosquito follows a negative binomial distribution with average oocyst number *M* and shape parameter *k*:

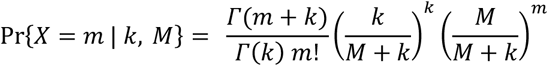

that gives the probability that the number *X* of oocysts in one mosquito is equal to *m*, for *m* = 0, 1, 2, etc. We determined the two parameters *M* and *k* using a Bayesian approach with the Metropolis-Hastings algorithm and determined their MLE (code available upon request). The estimates for *k* are given here:

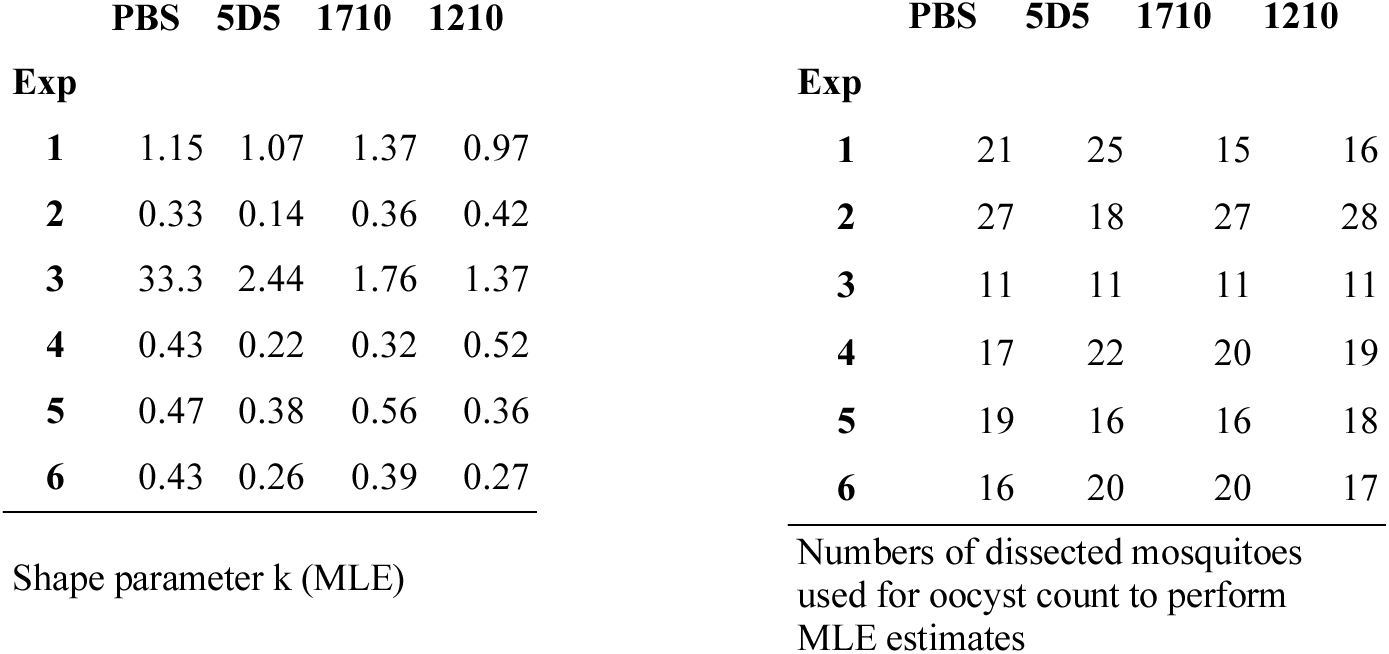

We then assumed that the number of sporozoites was linearly proportional to the number of oocysts (Vaughan et al., 1992; Stone et al., 2013; Miura et al., 2019). This allowed us to replace *M* (as derived from oocysts distribution) in the negative binomial with the average sporozoite number as given above. We used the MLE estimate of the shape parameter *k* and simulated 10,000 independent samples of mosquitoes of size given above. Each simulated sample is thus statistically identical to those provided by the experimental data.

Finally, for each two treatments and for each pair of simulated samples, we tested the null hypothesis that there is no difference between the treatments by random sampling while keeping sample sizes as in the experimental data. We thus created a distribution of the difference between the average number of sporozoites in the two treatments to be compared under this null hypothesis. The comparison of this distribution with the difference in sporozoite numbers as given by the experimental data produces the *p*-value listed below. The combined *p*-value using the Fisher method and true from false positives were discriminated using the Benjamini-Hochberg (BH) and the Benjamini-Liu (BL) methods at a false discovery rate *Q* = 10^-3^.

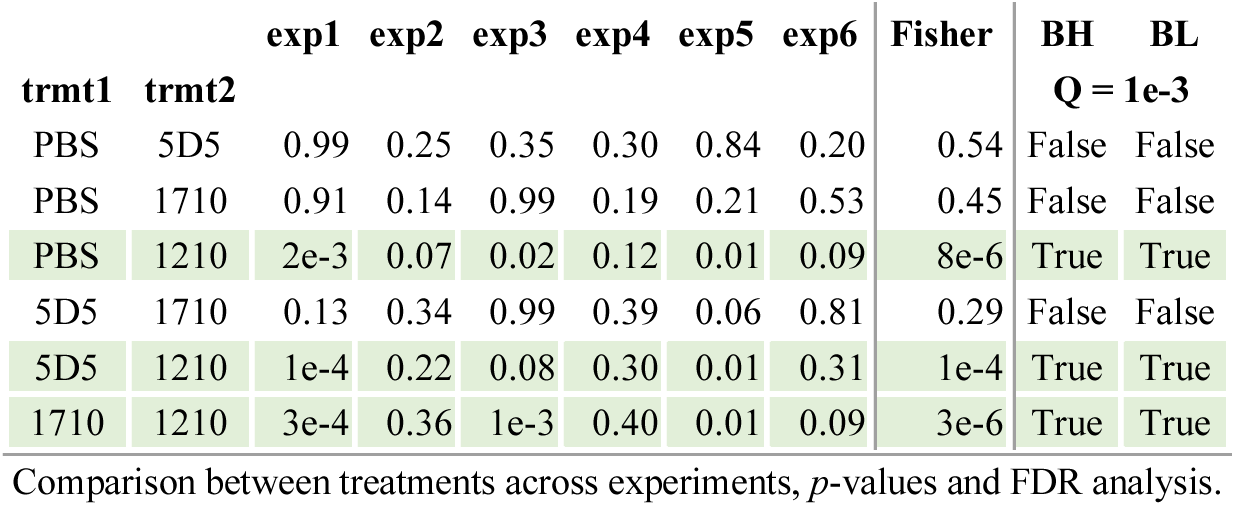

To check the consistency of this method, we simulated the sampled mosquito populations but skipped the shuffling across the samples.

### (Online) Supplemental Material

Supplemental materials include three tables and three figures. Supplemental tables: X-ray crystallography data collection and refinement (Table S1), description of primers used to generate mutant insert library for yeast display (Table S2), and PCR reactions and products for mutant insert library construction (Table S3). Supplemental figures: Experimental details of fluorescence-activated cell sorting of 5D5 IgG yeast display epitope mapping library and crystal structure (Figure S1), gating strategy for imaging flow cytometry quantification of mAb binding to live Pf sporozoites (Figure S2), and human mAbs against the PfCSP N-CSP identified from analysis of the PfSPZ-CVac samples (Figure S3).

## Author contributions

ET, GC, AW, RM, DO, SWS, IAW, HW, JPJ, EAL, conceived and designed the research; ET, GC, AW, RM, DO, NCW, KP, AB, NW, TP performed the research; ET, GC, AW, RM, DO, AV, SWS, IAW, NCW, PP, HW, JPJ, EAL, analyzed data; ET, GC, AW, HW, JPJ, EAL wrote the paper with input from all authors.

## Acknowledgements

We thank C. Kreschel for her support in Pf sporozoite production and cell culture, as well as H. Ahmed, L. Spohr, M. Andres and D. Eyemann (Vector Biology Unit, Max Planck Institute for Infection Biology, Berlin) for mosquito rearing and infections. The following reagents were obtained from BEI Resources, National Institute of Allergy and Infectious Diseases, National Institutes of Health: HC-04, hepatocyte (human), MRA-975, contributed by Jetsumon Sattabongkot Prachumsri. E.T. is currently supported by a CIHR Canada Graduate Scholarship and N.C.W. by NIH K99 AI139445. S.W.S was supported by a Hospital for Sick Children Lap-Chee Tsui Postdoctoral Fellowship and a Canadian Institutes of Health Research (CIHR) fellowship. This work was undertaken, in part, thanks to funding from the Bill and Melinda Gates Foundation (OPP1179906; J.-P.J, H.W. and E.A.L., and OPP1170236; I.A.W.), the CIFAR Azrieli Global Scholar program (J.-P.J.) and the Canada Research Chairs program (950-231604; J.-P.J.). X-ray diffraction experiments were performed using beamline 08ID-1 at the Canadian Light Source, which is supported by the Canada Foundation for Innovation, Natural Sciences and Engineering Research Council of Canada, the University of Saskatchewan, the Government of Saskatchewan, Western Economic Diversification Canada, the National Research Council Canada, and the Canadian Institutes of Health Research. The authors declare no competing financial interests.

## Notes

https://github.com/wchnicholas/CSP_Nterm_yeast_display

