## Supplementary Material for "A high-affinity antibody against the CSP N-terminal domain lacks *Plasmodium falciparum* inhibitory activity"

**Supplemental Tables and Figures.**

**Table S1. X-ray crystallography data collection and refinement data.**

|  | **5D5-CSP_81-98_** |
| --- | --- |
| **Wavelength (Å)** | 0.97959 |
| **Space group** | P2_1_ |
| **Cell dimensions** |  |
| ***a,b,c* (Å)** | 52.6, 60.9, 72.4 |
| **α, β, γ (°)** | 90, 97.7, 90 |
| **Resolution (Å)^a^** | 40-1.85 (1.95-1.85) |
| **No. molecules in ASU** | 1 |
| **No. observations** | 128,170 (18,612) |
| **No. unique observations** | 38,412 (5,588) |
| **Multiplicity** | 3.3 (3.3) |
| **R_merge_ (%)^b^** | 6.5 (45.0) |
| **R_pim_ (%)^c^** | 4.2 (29.0) |
| **<I/σ I>** | 13.7 (2.3) |
| **CC_½_** | 99.8 (73.7) |
| **Completeness (%)** | 98.7 (98.7) |
| **Refinement Statistics** |  |
| **Reflections used in refinement** | 38,396 |
| **Reflections used for R-free** | 1,920 |
| **Non-hydrogen atoms** |  |
| **5D5 Fab** | 3,341 |
| **PfCSP_81-98_** | 106 |
| **Solvent** | 420 |
| **R_work_^d^ /R_free_^e^** | 17.8 / 22.3 |
| **Rms deviations from ideality** |  |
| **Bond lengths (Å)** | 0.005 |
| **Bond angle (°)** | 0.80 |
| **Ramachandran plot** |  |
| **Favoured regions (%)** | 97.7 |
| **Allowed regions (%)** | 2.3 |
| **B-factors (Å^2^)** |  |
| **Wilson B-value** | 25 |
| **Average B-factors** | 33 |
| **5D5 Fab** | 32 |
| **PfCSP_81-98_** | 50 |
| **Solvent** | 38 |

a Values in parentheses refer to the highest resolution bin.

b Rmerge = Σhkl Σi | Ihkl, i - <Ihkl > | / Σhkl <Ihkl >

c Rpim = Σhkl [1/(N – 1)]1/2 Σi | Ihkl, i - <Ihkl > | / Σhkl <Ihkl >

d Rwork = (Σ | |Fo | − |Fc | |) / (Σ | |Fo |) - for all data except as indicated in footnote e.

e 5% of data were used for the Rfree calculation

**Table S2. Primers used to generate mutant insert library for yeast display.**

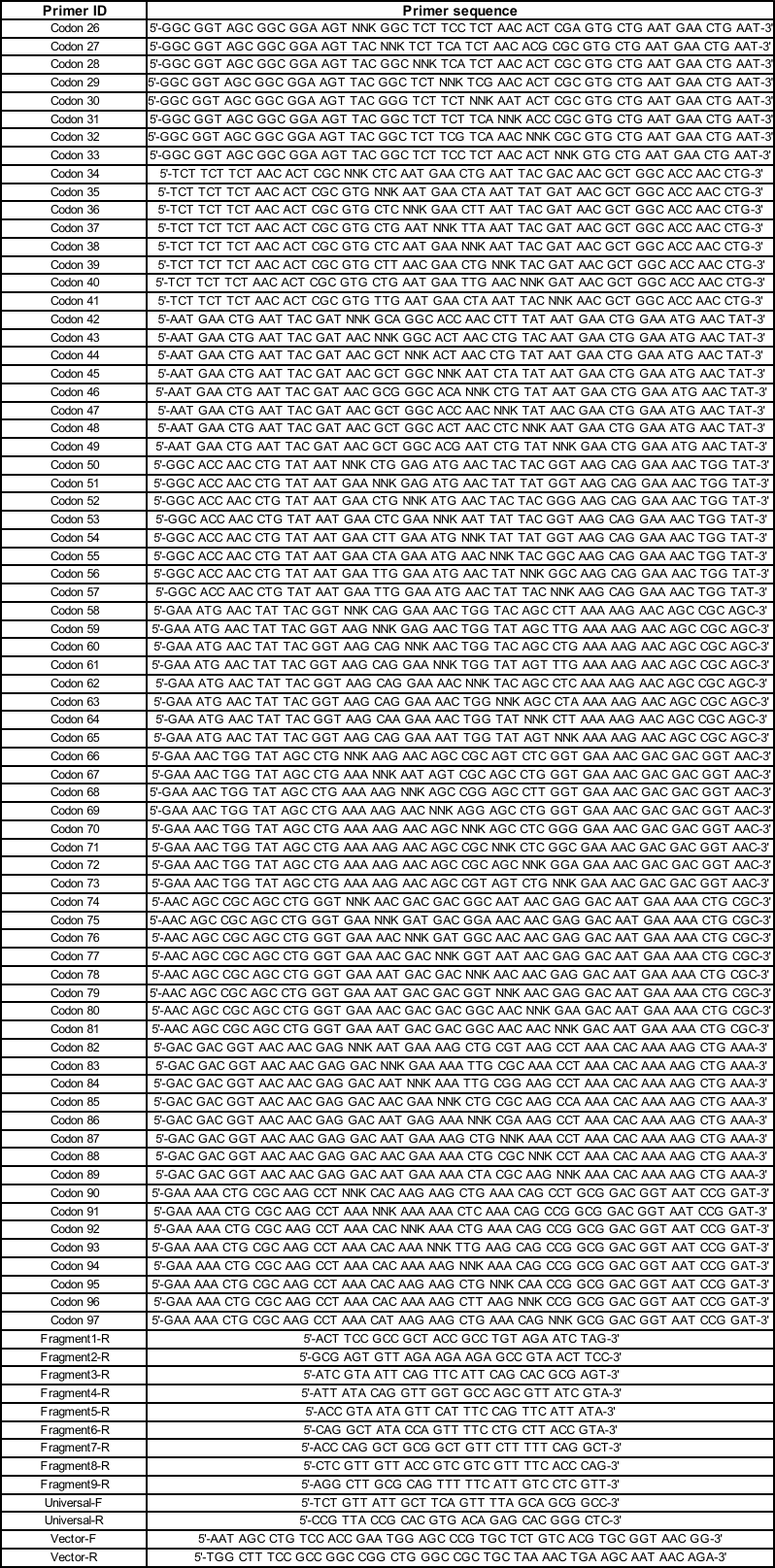

**Table S3. PCR reactions and products for mutant insert library construction**

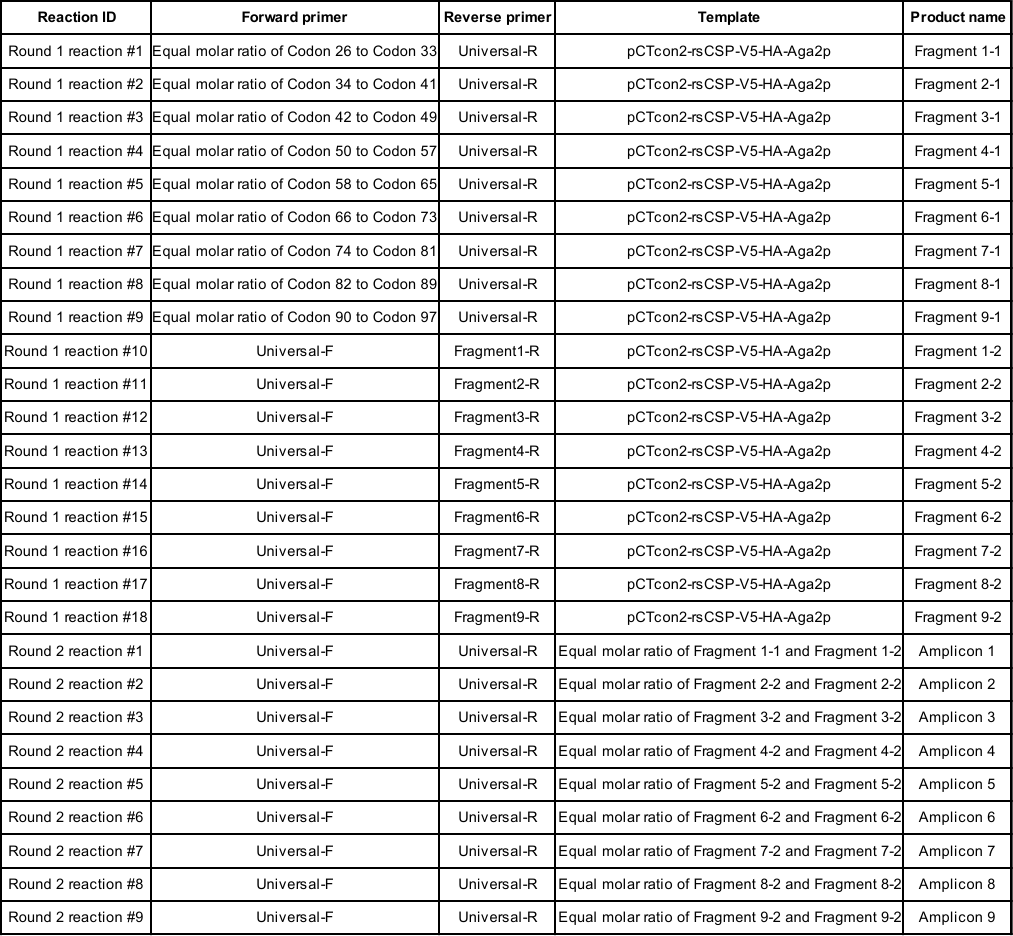

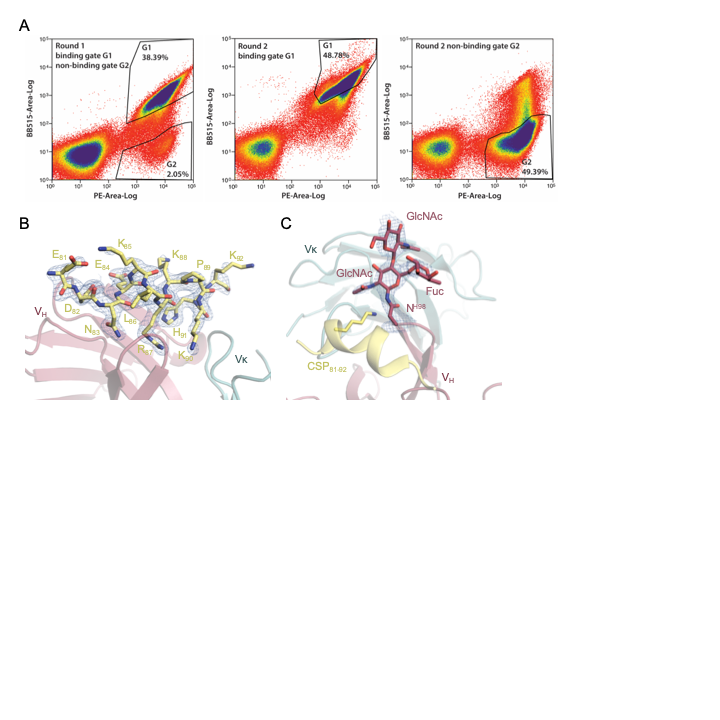

**Figure S1.** **Experimental details of fluorescence-activated cell sorting of 5D5 IgG yeast display epitope mapping library and crystal structure**. (A) Results from cell sorting (FACS) are shown. (Left) Round 1 sorting with 5D5 IgG binding (G1) and non-binding (G2) gates. (Middle) Round 2 sorting with 5D5 IgG binding gate. (Right) Round 2 sorting with 5D5 IgG non-binding gate. Percentages of cells are shown within the drawn gates. (B) Composite omit map electron density contoured at 1 sigma (blue mesh) around the N-CSP peptide and (C) the 5D5 N-linked glycan at position H.Asn98 of the HCDR3.

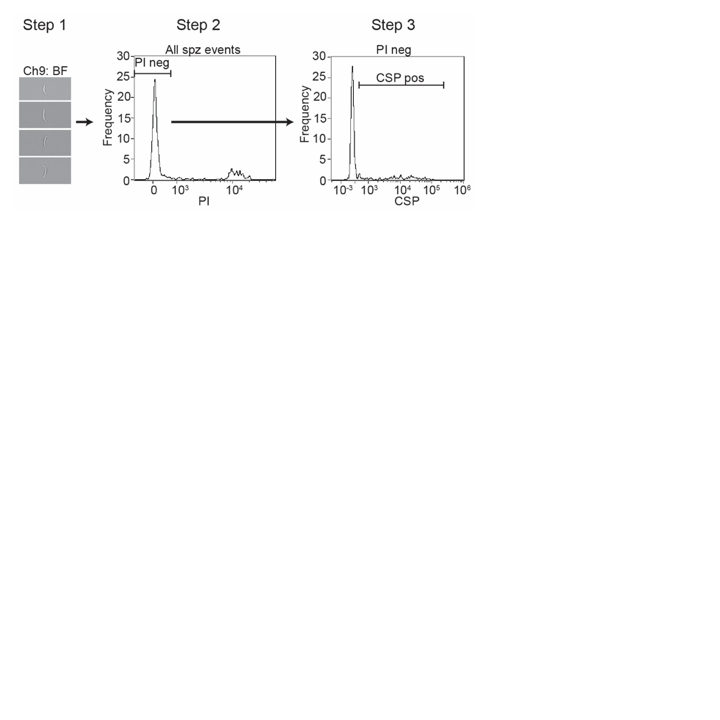

**Figure S2. Gating strategy for imaging flow cytometry quantification of mAb binding to live Pf sporozoites.** The gating strategy included 3 steps. Step 1: single in-focus sporozoites were manually selected on brightfield (BF) images. Step 2: live sporozoites were gated as the propidium iodide (PI)-negative population. Step 3: the proportion of mAb-positive sporozoites and MFI of mAb-bound sporozoites were quantified.

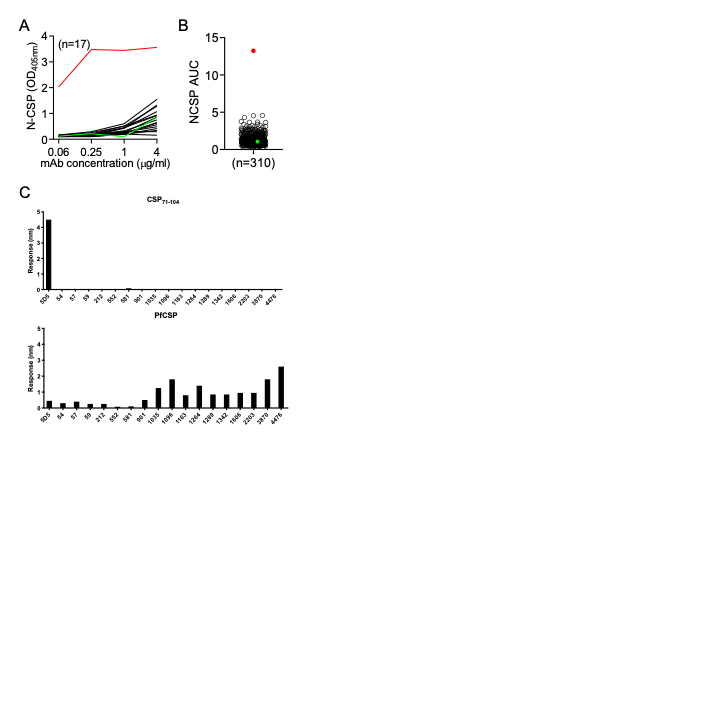

**Figure S3. Human mAbs against the PfCSP N-CSP were identified from analysis of the PfSPZ-CVac samples.** (A) Representative graph measuring binding of human mAbs isolated from memory B cells and plasmablasts (Murugan et al., 2018) to PfCSP N-CSP as measured in ELISA at the indicated antibody concentrations. Red and green line indicate 5D5 (positive control) and mGO53 (negative control), respectively. n indicates the number of antibodies shown. (B) Area under the ELISA curve (AUC) calculated from A for 310 human monoclonal antibodies. Red and green dots indicate 5D5 and mGO53, respectively. n indicates the number of antibodies tested. Data is representative of at least two independent measurements. (C) 17 human mAbs with an N-CSP AUC > 3 in at least one ELISA from B were tested for binding to CSP_71-104_ (top) and PfCSP (bottom) by BLI. mAbs were expressed as IgGs. 5D5 IgG was used as a positive control. Data is representative of two independent measurements.
